# Confirmatory evidence that miR-15a and miR-16 regulate BCL2 at the post-transcriptional level

**DOI:** 10.64898/2026.03.02.708996

**Authors:** Amelia Cimmino

## Abstract

The microRNAs miR-15a and miR-16 are key regulators of the anti-apoptotic oncogene BCL2, playing a significant role in tumorigenesis. These miRNAs function as tumor suppressors by directly targeting BCL2, whose overexpression contributes to cell survival and resistance to therapy in multiple malignancies, including chronic lymphocytic leukemia (CLL). The downregulation or deletion miR-15a/miR-16-1 cluster located on chromosome 13q occurs in about 50% of CLL patients and leads to the overexpression of the oncogenic BCL2, contributing to the survival and proliferation of cancer cells. In this confirmatory study, we provide additional evidence supporting the mechanism by which these miRNAs mediate the inhibition of BCL2 translation, leading to reduced levels of BCL2 protein with no significant effect on BCL2 mRNA degradation. This mechanism has been previously established as a critical pathway in the regulation of apoptosis, particularly in cancer cells where BCL2 overexpression is often associated with resistance to cell death. Our findings reinforce the notion that miRNAs, such as miR-15 and miR-16, bind to the 3’-UTR of BCL2 messenger RNA (mRNA), specifically repressing its translation without inducing mRNA degradation. The results from our study align with previous research, confirming that the miRNA-mediated inhibition of BCL2 translation serves as a precise regulatory mechanism that targets protein synthesis rather than mRNA stability. These findings highlight the role of miRNAs in fine-tuning post-transcriptional gene regulation, offering a targeted approach to downregulate oncogenic proteins like BCL2 without disrupting the underlying mRNA, which could be leveraged for more refined therapeutic strategies.

## Introduction

MicroRNA (miRNAs) are an abundant class of 19–22 nucleotide (nt) long noncoding RNAs that regulate gene expression. The mechanism by which miRNAs inhibit the translation of their target genes primarily involves post-transcriptional gene regulation. Based on sequence complementarity, a short (5–7nt long) sequence in the miRNA, identified as the seed sequence, determines the target messenger RNA (mRNA) binding specificity, typically in the 3’untranslated region (3’UTR) region of the mRNA. This interaction leads to two main outcomes: translation inhibition or mRNA degradation, depending on the degree of complementarity and the cellular context. MiRNAs either degrade the targeted mRNAs or suppress their translation. One well-characterized miRNA family is the miR-15 family, consisting of the three bicistronic clusters in humans encoding for miR-15a/16-1 (on chromosome 13q), miR15b/16-2 (on chromosome 3p) and miR-497/195 (on chromosome 17p), with miR-15a/16-1 best known for its significant tumor suppressive impact in chronic lymphocytic leukemia (CLL) [1].

The first focus of miR-15a/16-1 research was on its function in controlling the apoptotic pathway [2]. Hsa-miR-15a/16-1 and hsa-miR-15b/16-2 clusters exhibit high levels of expression during B cell maturation, while miR-497 and miR-195 are mostly expressed in non-immune cells [3]. The chromosomal region encoding the miR-15a/16-1 cluster is lost in approximately 60% of B-cell chronic lymphocytic leukemia (CLL) patients, a disease for which the miR-15a/16-1 is well known for its tumor-suppressive properties [4]. Their relevance has been confirmed in mice models where the deletion of the miR-15a/16-1 and miR-15b/16-2 clusters generated a CLL-like disease [5,6]. Using computational miRNA target prediction (targetscan.org), in our previous study we found that miR-15a and miR-16-1 target the 3′UTR of BCL2 mRNA [2].

Here, we provide additional confirmation of the mechanism of miRNA-mediated inhibition of translation of BCL2 that reduces the levels of BCL2 protein but not the BCL2 mRNA levels.

## Materials and Methods

### Cell lines and cell culture

The human megakaryocytic MEG-01 cell line was derived from the bone marrow of a 55-year-old patient with Chronic Myelogenous Leukemia. The cell line carries the fusion gene BCR/ABL resulting from the Philadelphia chromosome balanced translocation t(9;22). MEG-01 cells were maintained in RPMI-1640 medium modified to contain 2 mM L-glutamine, 4500 mg/L glucose, 10% of fetal bovine serum and 1% penicillin and streptomycin.

### Transfection Assays

MEG-01 was grown in 10% FBS in RPMI medium 1640, supplemented with 2 mM L-glutamine, 4500 mg/L glucose, 10% of fetal bovine serum, 1 nonessential amino acid and 1 mmol sodium pyruvate at 37°C in a humified atmosphere of 5% CO2. The cells were cotransfected in 12-well plates by using Lipofectamine RNAiMAX (Invitrogen). For each well 50 nM miR-16-1-sense and miR-15a-sense and anti-miR-16-1 and anti-mir-15a precursor miRNA inhibitor (Thermo Fisher Scientific) were used.

### Gene expression analysis

Total RNA was extracted using TRIzol® reagent (Thermo Fisher Scientific), and cDNA was synthesized using the SuperScript III First-Strand Synthesis System (Thermo Fisher Scientific). BCL2 mRNA expression was analyzed by semi-quantitative PCR and RT-qPCR. Semi-quantitative PCR was performed using the Titan One Tube RT-PCR system according to the manufacturer’s instructions (Roche). The primers used for BCL2 amplification were forward (FW) 5′-GTGGCCTTCTTTGAGTTCGG-3′ and reverse (RV) 5′-ACCAGGGCCAAACTGAGCA-3′. Actin was used as an internal control with the following primers: FW 5′-CCCATGCCATCCTGCGTCT-3′ and RV 5′-GAAAGGGTGTAACGCAACTAA-3′. For RT-qPCR analysis, the same primer pairs were used for BCL2, while actin was amplified using the following primers: FW 5′-ATCTGAGGAGGGAAGGGGAC-3′ and RV 5′-AGACCTGTACGCCAACACAG-3′. Relative gene expression was calculated using the 2^(-ΔΔCt) method, where ΔCt represents the difference between the Ct values of the target gene and the reference gene, and ΔΔCt represents the difference between experimental and control samples.

### MiRNA expression analysis

MiRNA expression levels were measured by RT-qPCR using TaqMan MicroRNA Assays (Applied Biosystems) specific for mature miR-15a (Assay ID 477858) and miR-16-1 (Assay ID 477860). Amplification and detection were performed using TaqMan Universal PCR Master Mix. Relative miRNA expression was calculated using the 2^(-ΔΔCt) method, with U6 small nuclear RNA used as the endogenous reference. Statistical comparisons were performed using unpaired t-tests in GraphPad Prism software (version 10.3.1).

### Western blot analysis of BCL2 protein expression

Following transfection, MEG-01 cells were harvested and lysed in ice-cold lysis buffer supplemented with protease inhibitors. Total protein concentration was determined using a standard protein assay. Equal amounts of protein were separated by SDS-PAGE and transferred onto PVDF membranes using standard Western blotting procedures. Membranes were blocked and incubated with a mouse monoclonal anti-BCL2 antibody (Santa Cruz Biotechnology) overnight at 4°C. After incubation with the appropriate HRP-conjugated secondary antibody, immunoreactive bands were visualized using an enhanced chemiluminescence detection system. Protein loading and transfer efficiency were assessed by reprobing the membranes with a mouse monoclonal anti-GAPDH antibody (Cell Signaling), which was used as an internal loading control.

## Results

### Evaluation of miR-15a/miR-16-1 transfection efficiency

MEG-01 cells were transfected with miR-15a and miR-16-1 sense (mimic) or antisense (AS) oligonucleotides, either individually or in combination, at a final concentration of 50 nM. Transfection efficiency was assessed by measuring miR-16-1 expression using RT-qPCR. As shown in **Figure 1A**, cells transfected with miR-15a/16-1 mimics or miR-16-1 mimic alone displayed a robust increase in miR-16-1 expression, confirming effective miRNA delivery. Conversely, transfection with anti-miRNAs (miR-15a+16-1_AS or miR-16-1_AS) resulted in a significant reduction of endogenous miR-16-1 levels confirming efficient modulation of miRNA expression.

**Figure 1.**
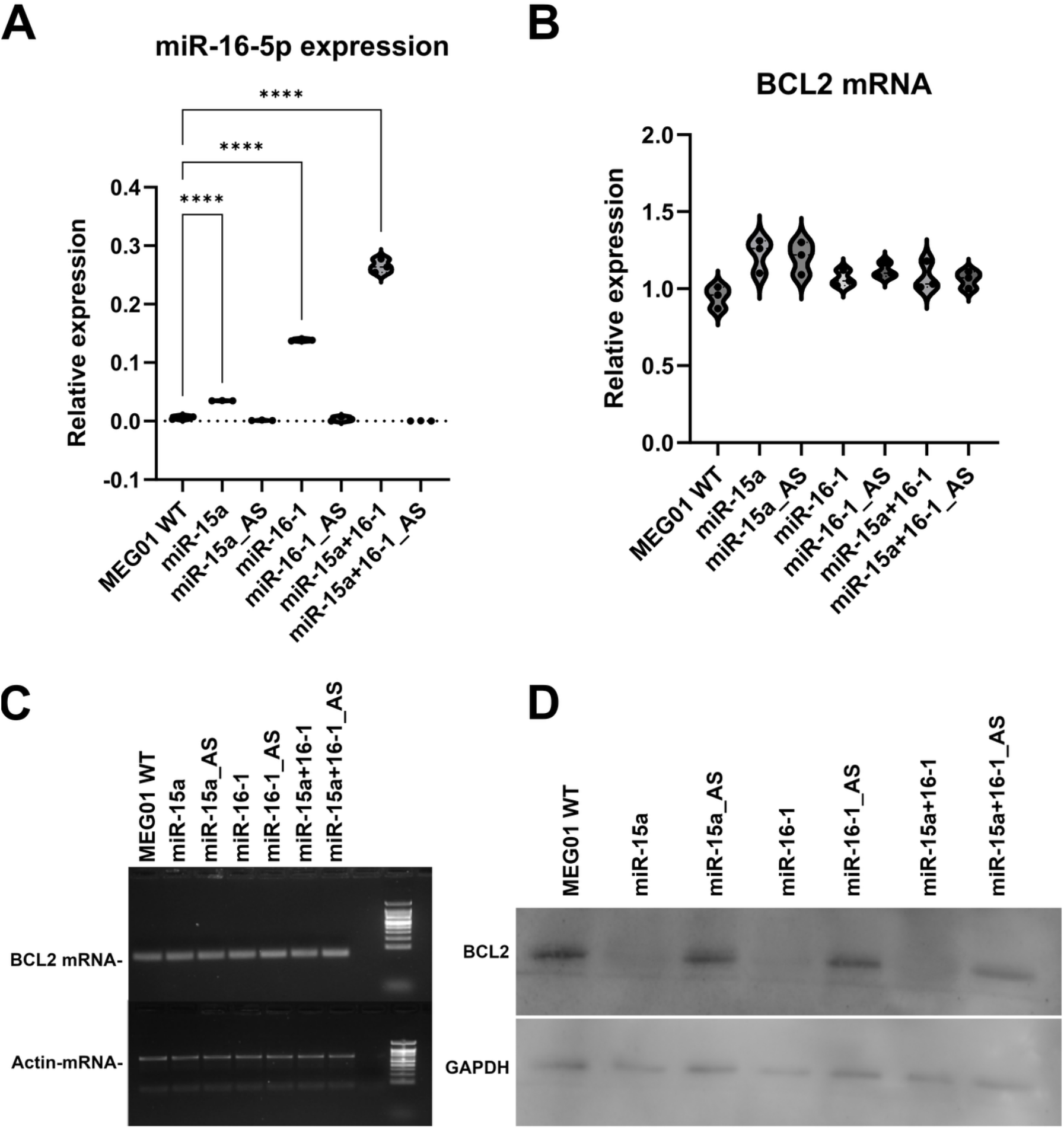
Transfection of the miR-15a/miR-16-1 cluster in MEG-01 cells. (A) Relative expression of miR-16-5p measured by RT-qPCR in MEG-01 wild-type (WT) cells transfected with miR-15a or miR-16-1 mimics, corresponding antisense oligonucleotides (AS), or their combinations. Data represent three independent experiments. **** *p* < 0.001. (B) Relative BCL2 mRNA expression in the same experimental conditions, determined by RT-qPCR using the 2^−ΔCt formula and actin as the reference gene. (C) Semi-quantitative RT-PCR analysis of BCL2 mRNA levels in MEG-01 cells under the indicated conditions, normalized to actin expression. (D) Western blot analysis of BCL2 protein expression in MEG-01 cells, with GAPDH used as a loading control.

### Evaluation of BCL2 mRNA expression following miR-15a/miR-16-1 modulation

To determine whether miR-15a/miR-16-1 modulates BCL2 expression at the mRNA level, BCL2 transcript levels were analyzed by RT-qPCR (**Figure 1B**) and semi-quantitative RT-PCR (**Figure 1C**). No significant differences in BCL2 mRNA levels were detected between transfected cells and MEG-01 wild-type controls (**Figures 1B,C**), indicating that miR-15a and miR-16-1 do not affect BCL2 mRNA stability and suggesting that BCL2 regulation does not occur through mRNA degradation.

### Post-transcriptional regulation of BCL2 protein expression by miR-15a and miR-16

BCL2 has been previously identified as a direct target of post-transcriptional repression by miR-15a and miR-16-1 [2]. In our earlier study, luciferase reporter assays demonstrated a direct interaction between miR-15a/miR-16-1 and the BCL2 3′UTR, confirming sequence-specific targeting (2). Consistently, Western blot analysis in the same work showed that both miR-15a and miR-16-1 significantly downregulate BCL2 protein expression (2). Since miRNAs can modulate gene expression by affecting either mRNA stability or translational efficiency (19), we investigated here the molecular level at which the miR-15a/16-1 cluster regulates BCL2 expression in MEG-01 cells.

Western blot analysis revealed a marked reduction in BCL2 protein levels in cells transfected with miR-15a or with miR-16-1 mimics, with a stronger effect observed upon combined transfection (**Figure 1D**). This repression was reversed by transfection with the corresponding antisense oligonucleotides. The clear dissociation between stable BCL2 mRNA levels and reduced protein expression provides strong evidence that the miR-15a/miR-16-1 cluster predominantly represses BCL2 at the post-transcriptional level, most likely through inhibition of mRNA translation rather than mRNA degradation.

## Discussion

MicroRNAs with incomplete target complementarity can promote target repression by at least two different mechanisms, resulting either in mRNA degradation or inhibition of mRNA translation. In the first scenario, miRNAs, rather than affecting mRNA stability, inhibit mRNA translation at the step of initiation [4–6] or elongation [7]. Having already established the ability of miR-15a-16-1 to regulate BCL2 protein levels by western blot and luciferase assay, here we re-assayed its effect on BCL2 protein expression. The specific interaction between the miRNA seed region and the BCL2 mRNA is essential for the translational repression mechanism. This interaction does not activate mRNA decay pathways but instead results in efficient inhibition of protein synthesis. Accordingly, several independent studies have reported a significant downregulation of BCL2 protein levels following miR-15a/miR-16-1 expression, while BCL2 mRNA levels remain unchanged, as demonstrated by combined qRT-PCR and Western blot analyses [8,9]. This miRNA-mediated translational repression of BCL2, occurring without detectable mRNA degradation, underscores the precision of post-transcriptional gene regulation and highlights a mechanism that is particularly relevant in therapeutic contexts where reduction of protein levels without altering mRNA stability is desirable.

